# Spatial-proteomics reveal *in-vivo* phospho-signaling dynamics at subcellular resolution

**DOI:** 10.1101/2021.02.02.425898

**Authors:** Ana Martinez-Val, Dorte B. Bekker-Jensen, Sophia Steigerwald, Adi Mehta, Trung Tran, Krzysztof Sikorski, Estefanía Torres-Vega, Ewa Kwasniewicz, Sólveig Hlín Brynjólfsdóttir, Lisa B. Frankel, Rasmus Kjøbsted, Nicolai Krogh, Alicia Lundby, Simon Bekker-Jensen, Fridtjof Lund-Johansen, Jesper V. Olsen

**Affiliations:** Novo Nordisk Foundation Center for Protein Research, Proteomics Program, Faculty of Health and Medical Sciences, University of Copenhagen, Copenhagen, Denmark; Evosep Systems, Odense, Denmark; Max Planck Institute of Biochemistry, Department of Proteomics and Signal Transduction, Martinsried, Germany; Department of Immunology, Oslo University Hospital, Rikshospitalet, Postboks 4950, Nydalen, 0424 Oslo, Norway; Cardiac Proteomics, Faculty of Health and Medical Sciences, University of Copenhagen, Copenhagen, Denmark; Center for Healthy Aging, Department of Cellular and Molecular Medicine, University of Copenhagen, Copenhagen, Denmark; Danish Cancer Society, Copenhagen, Denmark; Department of Nutrition, Exercise and Sports, University of Copenhagen, Copenhagen, Denmark; Department of Cellular and Molecular Medicine, University of Copenhagen, Copenhagen, Denmark

## Abstract

Dynamic change in subcellular localization of signaling proteins is a general concept that eukaryotic cells evolved for eliciting a coordinated response to stimuli. Mass spectrometry (MS)-based proteomics in combination with subcellular fractionation can provide comprehensive maps of spatio-temporal regulation of cells, but involves laborious workflows that does not cover the phospho-proteome level. Here we present a high-throughput workflow based on sequential cell fractionation to profile the global proteome and phospho-proteome dynamics across six distinct subcellular fractions. We benchmarked the workflow by studying spatio-temporal EGFR phospho-signaling dynamics in-vitro in HeLa cells and in-vivo in mouse tissues. Finally, we investigated the spatio-temporal stress signaling, revealing cellular relocation of ribosomal proteins in response to hypertonicity and muscle contraction. Proteomics data generated in this study can be explored through https://SpatialProteoDynamics.github.io.

---

Protein function is tightly controlled in cells through multiple mechanisms. Protein activity can be dynamically modulated, for instance, by changing translation rate ^1^ or by post-translational modifications, such as site-specific phosphorylation or ubiquitination ^2^. Moreover, most proteins do not operate in isolation, but rather they need to interact with other proteins to elicit their functions ^3^. Most of these regulatory mechanisms have been the subject of extensive research. Additionally, a protein’s function can also be regulated in a spatial manner by modulating its subcellular localization. This regulatory layer, *i.e.* cellular compartmentalization, is especially important for faithful transmission through signal transduction pathways, where fast responses are required, such as nucleocytoplasmic shuttling of transcription factors for transcriptional control ^4^ or endocytic internalization of activated receptors for degradation or recycling ^5^. Moreover, it is well established that many proteins can exert different functions depending on their subcellular location ^6^. Due to its biological importance, subcellular localization of proteins has been studied extensively, mainly by using molecular biology techniques, relying on either imaging ^7^, or, most recently, on information derived from proximity-labeling experiments ^8^. Although very sensitive and powerful, these techniques lack throughput as they cannot provide information on protein location at a proteome-wide level. In recent years, several studies presented the potential of MS-based proteomics to explore the subcellular proteome. Among them, approaches such as LOPIT-DC ^9^ or SubCellBarcode ^10^ stand out due to their sensitivity, coverage and resolution, allowing to map the location of more than 8000 proteins. However, both methods rely on isobaric tandem mass tag labeling for accurately quantifying subcellular protein localization, and they require extensive off-line peptide fractionation and consequently lengthy liquid chromatography coupled to mass spectrometry (LC-MS/MS) analysis time to achieve the desired depth on the proteome, thus minimizing throughput. Conversely, other studies have proposed single-shot LC-MS/MS analysis and label-free quantification as an alternative to obtain faster organellar-maps ^11,12^ at the expense of coverage allowing to map the location of about 4,000 proteins in ~12 hours of MS analysis. Consequently, there is still a gap of knowledge in how to obtain deep subcellular proteomes without compromising MS-time to enable the application of these techniques to different experimental conditions with multiple biological replicates.

Subcellular translocation of a protein is a dynamic regulatory event, and it is therefore essential to incorporate the temporal dimension when studying protein translocation in response to stimuli. Spatio-temporal proteomics is very challenging, and only few studies have been performed, such as the spatio-temporal characterization of cytomegalovirus infection ^13^. This is probably due to the complexity in applying these workflows in a high throughput manner, which substantially limits their usefulness to analyze global subcellular proteome dynamics. Moreover, to our knowledge, none of the current approaches to study the subcellular proteome changes have covered the signaling layer of the phospho-proteome, which is known to control and trigger protein relocation ^14^. To overcome these limitations, and provide an accessible workflow to study spatio-temporal phospho-proteome regulation, we present a workflow based on a sequential subcellular fractionation protocol that when coupled to fast chromatographic LC-MS/MS analysis using data-independent acquisition (DIA) provides rapid, sensitive and reproducible subcellular phospho-proteome maps. Importantly, our high-throughput approach allows studying spatial-dynamics in a temporal manner in both cell lines and tissues with multiple replicates. To demonstrate the general applicability of the spatial proteomics workflow, we have applied our method in two different biological settings to study the spatio-temporal response of the proteome and phospho-proteome both *in vitro* and *in vivo*.

## Results

### High-throughput and reproducible maps of subcellular phospho-proteomes using directDIA-MS spatial proteomics

Chemical fractionation provides an attractive alternative to more elaborate methods for separation of intact organelles. However, reproducibility is hampered by the widespread use of a plethora of kits with undisclosed composition and the lack of meta-analysis ^15^. Here, we present a streamlined pipeline for the analysis of (phospho)-proteome dynamics in distinct subcellular compartments of the cell (Fig. 1A). Six extraction buffers with different detergent, salt and chemical composition (see Methods) ^16^, were used sequentially to profile six distinct subcellular fractions in cell lines and tissues at both proteome and phospho-proteome level. In contrast to already published MS-based subcellular fractionation methods, our approach can be employed in a high-throughput manner. Due to the use of directDIA and fast chromatographic gradients, quantitative data for subcellular proteomes and phospho-proteomes can be generated for six subcellular fractions in just 5 hours of MS time. This enables the possibility to include multiple biological replicates as well as different experimental conditions or time points.

**Figure 1:**
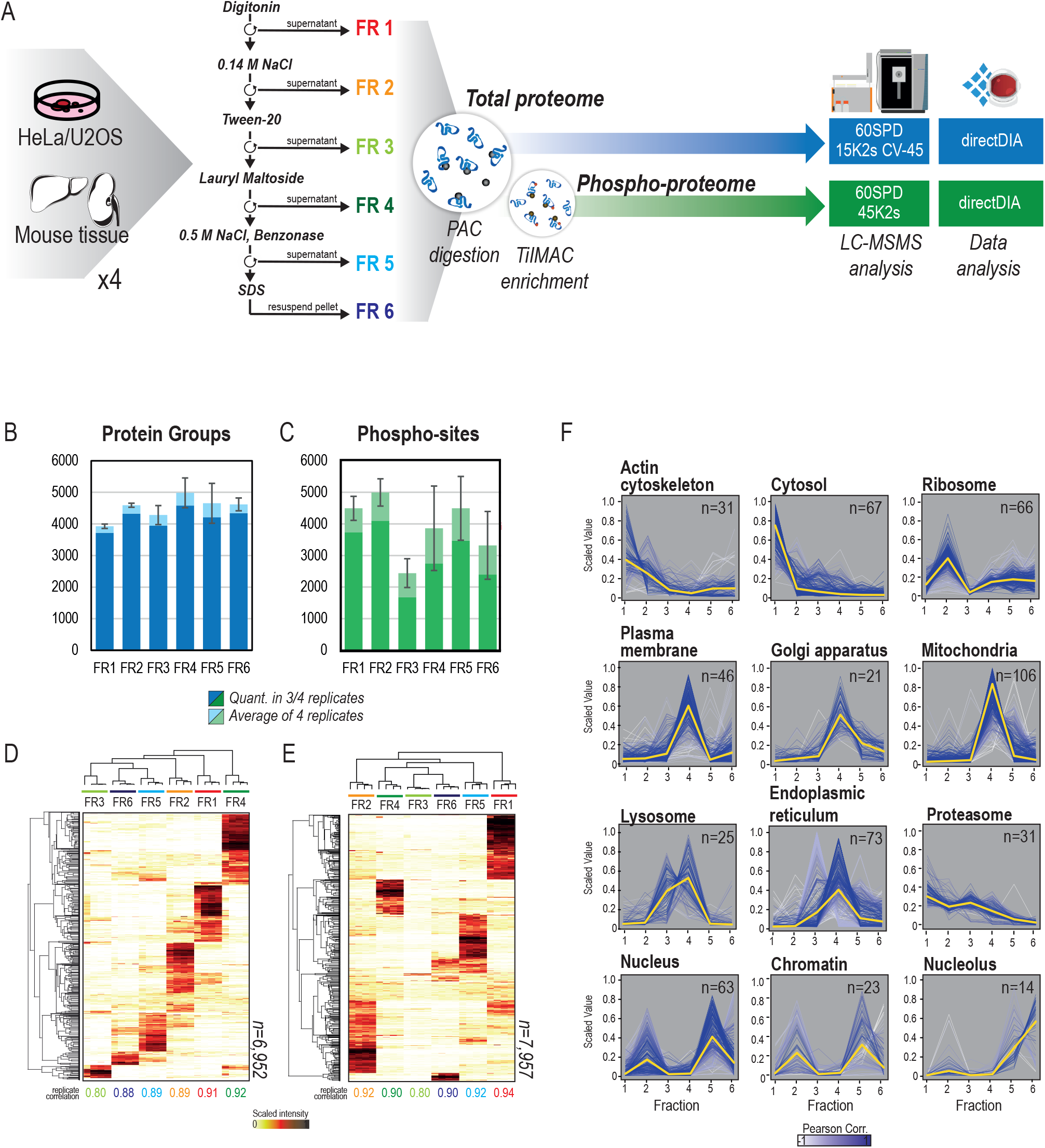
High-throughput and reproducible subcellular fractionation. (A) Experimental workflow for subcellular fractionation and LC-MS data acquisition. (B) Bar-plot summary of the identified proteins as average of 4 replicates per fraction (light blue bar) and quantified proteins in at least 3 replicates (dark blue). The error bars indicate the standard deviation in identification number between replicates. (C) Bar-plot summary of the identified phosphorylation sites as average of 4 replicates per fraction (light green) and quantified phosphorylation sites in at least 3 replicates (dark green). The error bars indicate the standard deviation in identification number between replicates. (D) Heatmap of scaled intensities per replicate, of four replicates, of the subcellular proteome, showing both protein and sample clustering. (E) Heatmap of scaled intensities per replicate, of four replicates, of the subcellular phospho-proteome, showing both protein and sample clustering. (F) Profile-plots of cell compartment markers in the subcellular proteome HeLa dataset. Scaled intensity across fractions is plotted for each independent replicate. Gradient of white to blue indicates Pearson correlation to the centroid of each distribution, which is highlighted as a yellow line.

To evaluate the coverage and reproducibility of the workflow, we applied our method to HeLa human cervix carcinoma cells. Four biological replicates of HeLa cells were serum starved overnight, and trypsinized just prior subcellular fractionation. The entire experiment was performed at 4°C in the presence of protease and phosphatase inhibitors. Once the fractions were collected, they were lysed in boiling SDS and reduced/alkylated before being subjected to protein aggregation capture (PAC)-based tryptic digestion ^17^ using the fully automated KingFisher platform. Approximately five percent of the resulting peptide mixtures from each subcellular fraction were used for total proteome analysis. Remaining peptides were subjected to phosphopeptide enrichment using TiIMAC-HP beads, also on the KingFisher platform. To increase measurement depth of our single-shot subcellular proteomes, we took advantage of high field asymmetric waveform ion mobility spectrometry (FAIMS) in combination with fast scanning DIA acquisition methods ^18^ while employing a more sensitive MS-acquisition method for phospho-proteomics samples (Fig. 1A). In both cases, we used a library-free approach (directDIA) in Spectronaut to analyze the resulting raw MS data. In total, we could quantify 6952 proteins and 7957 phosphorylation-sites in our dataset. On average, 4000 proteins were identified in each fraction and more than 90% of them were robustly quantified in three out of four replicates (Fig. 1B). In contrast, the phospho-proteome profile of each fraction yielded more variable results per fraction (Fig. 1C), where fractions 3 and 6 reproducibly result in less identified phosphorylation sites, which can be due to intrinsic lack of phosphorylation events in those specific subcellular compartments.

Since the current protocol is based on the sequential fractionation of cellular compartments, we hypothesized that the MS signal intensity for a given protein measured per fraction should represent its relative abundance in the whole cell proteome. To evaluate this, we plotted the sum of each protein intensity across fractions against the average intensity observed in a total lysate, and found a good correlation (Spearman correlation=0.7, p-value: 4e-324) between both datasets, confirming our expectations (Suppl. Fig. 1A).

Hierarchical clustering analysis of the scaled fractional intensities of proteins and phosphorylation sites reveals high reproducibility between experimental replicates with Pearson correlation between replicates in the range 0.80-0.94 (Fig. 1D-E, Suppl. Fig 1B.). Most importantly, the intensity profiles for both proteins and phosphorylation sites across fractions reveal very well-defined clusters corresponding to the proteins enriched in each cellular compartment purified in each fraction, reflecting the resolving power of the fractionation method (Fig. 1D-E). To define which cell compartment was enriched in each fraction, we annotated the dataset using established subcellular protein markers ^19,20^ and plotted their profile distribution across fractions at the proteome level (Fig. 1F). This analysis revealed three major cellular compartments in our dataset: fractions 1-2 corresponding to cytosolic and cytoskeletal proteins, fractions 3-4 to plasma membrane and membranous organelles, and fractions 5-6 to the nucleus (Fig. 1F). Moreover, we also observed a very clear distinction of nuclear components between fractions 5 and 6, with nucleoplasm and chromatin-bound proteins mainly purified in fraction 5, whereas nucleolar proteins are mostly observed in fraction 6 (Fig. 1F). Gene Ontology (GO) enrichment analysis in each protein cluster revealed more specific patterns for each fraction. Interestingly, we observed that many protein complexes tend to be purified in fraction 2, including ribosomes, the exocyst or the septin complex. On the other hand, many proteins involved in cell cycle and DNA replication are co-purified in fraction 2, possibly due to cells undergoing cell division, in which the nuclear membrane is dissolved. To validate the subcellular localization of the proteins identified, we overlapped the dataset with the subcellular annotations from the more comprehensive imaging-based Cell Atlas ^7^ and found that the subcellular patterns were reproduced (Suppl. Fig. 2A). Reassuringly, the observed subcellular protein profiles were highly reproduced at the phospho-proteome level (Suppl. Fig. 2B), with few interesting variations. For instance, whilst ribosomal proteins are highly enriched in fraction 2 at the proteome level, the phosphorylated counterparts of the same ribosomal proteins are also significantly enriched in fraction 1. This observation could indicate an intrinsic difference in compartmentalization of ribosomal proteins depending on their phosphorylation status, but this needs further evidence.

Next, we investigated the relationship between subcellular localization and global phosphorylation events. To do so, we plotted the profiles of 221 kinases identified in our dataset (Suppl. Fig. 2C). Interestingly, we found that fraction 3 contained less kinases compared to the other fractions, which aligns well with our previous observation of lower phosphorylation events reported in this specific compartment (Fig. 1C). Furthermore, to confirm that both a kinase and its substrate were localized in the same cellular compartment, we evaluated some well-known kinase substrate relationships and found that they generally co-localize (Suppl. Fig. 2D). Finally, we assessed the reproducibility of the subcellular protein localization between different cell lines by repeating the analysis in U2OS human osteosarcoma cells, and found the fractionation resolution to be highly reproducible indicating conserved subcellular proteome distribution across the two cell lines (Suppl. Fig. 2E). Profiling the known subcellular protein markers in U2OS cells, and comparing the correlation of their profiles against those obtained for HeLa (Suppl. Fig. 2F-G), also showed good reproducibility of the technique independently of the two cell lines.

To benchmark our results against previously published datasets, we used the MetaMass ^15^ tools for meta-analysis of subcellular proteomics data (see Methods) ^15^. MetaMass analyses a list of gene names and assigned groups obtained by k-means clustering of normalized MS data and compares this to several built-in sets of markers for subcellular compartments. The output includes statistics for precision and recall for the markers as well as the harmonic mean of the two (F-score). The markers used here correspond to subcellular locations mapped for U2OS cells by MS analysis of fractions obtained by organelle separation ^7^. We compared our dataset against a reference published dataset obtained by chemical fractionation of KM12 colorectal carcinoma cells using a widely used commercial fractionation kit ^21^ (Fig 2A-B). The method presented here yielded higher F-scores for partitioning of the identified marker proteins from HeLa cervical carcinoma cells and U2OS osteosarcoma cells into the correct subcellular compartments than those calculated for the reference dataset (Fig. 2C). An unexpected finding was that the F-scores for markers of the plasma membrane, endoplasmic reticulum, lysosomes and mitochondria were in the range of 0.69-0.85 (Fig 2C). Similarly, our method reported well-defined fractions and comparable results as other published methods, such as differential density centrifugation ^7,9,22^ or differential centrifugation ^9,10^ (Suppl. Fig 3A-E), whilst minimizing input requirements, simplifying sample preparation and reducing MS acquisition time (Suppl. Table 1). While the precision obtained with centrifugation-based methods for organelle separation is higher as expected, it seems that chemical fractionation can yield approximate locations.

**Figure 2:**
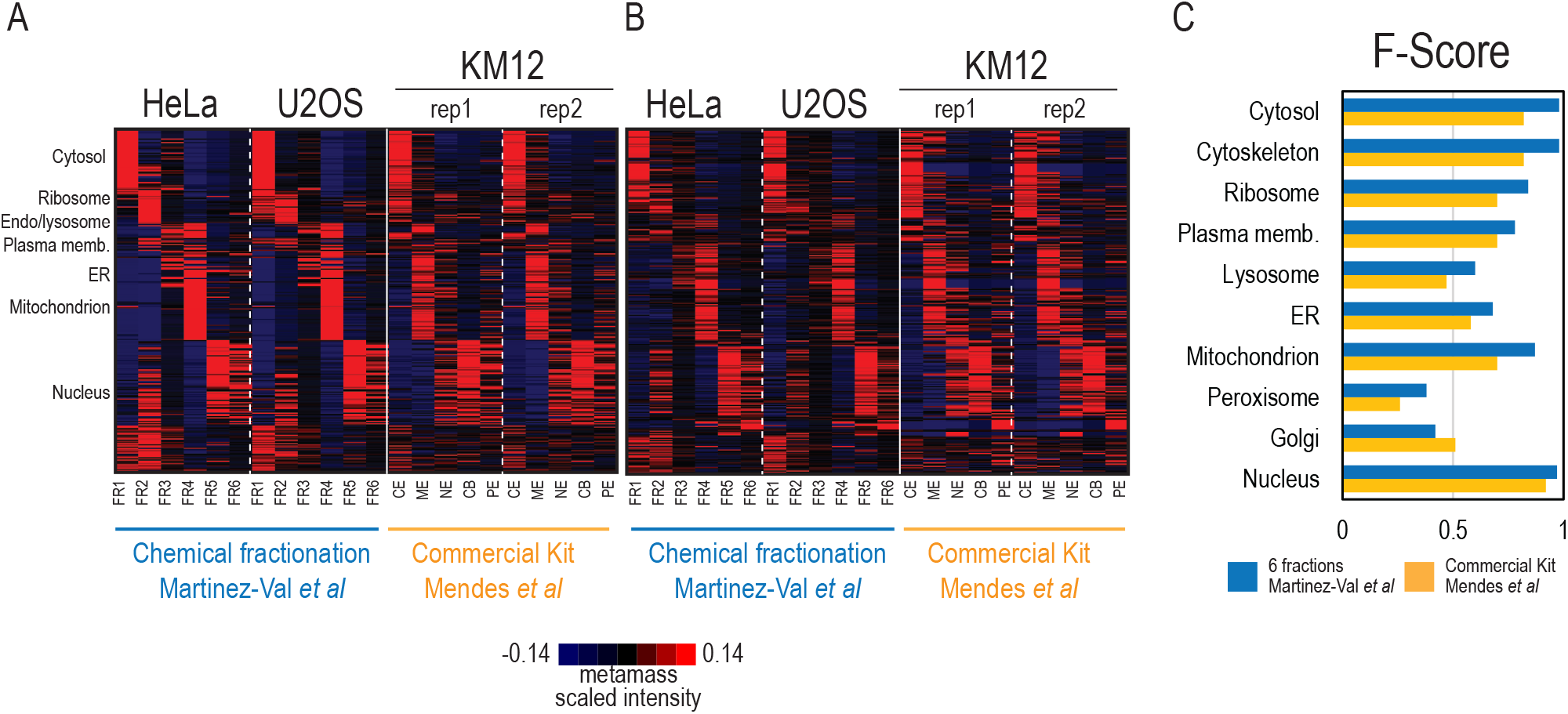
Comparative analysis of subcellular fractionation protocols based on chemical reagents using Metamass. (A-B) Heatmaps showing protein distribution across fractions obtained from HeLa and U2OS using the present subcellular fractionation protocol and KM12 using the commercial kit (Pierce) employed in Mendes et al (PMID: 28861940). Proteins were classified and sorted using the Excel-based analysis tool MetaMass. In (A) proteins are clustered based on the data from HeLa and U2OS fractionation in the present study. In (B) proteins are clustered based on Mendes *et al* data. Heatmaps were obtained after normalizing gene distribution and center samples by mean in Cluster 3.0, and plotted in TreeView. CE: cytoplasmic extract. ME: membrane extract. NE: nuclear extract. CB: chromatin-bound. PE: pellet extract. ER: Endoplasmic reticulum. (C) F-score barplot for the protein assignment to organelles in the present study (blue) and in Mendes *et al* (yellow).

### Spatio-temporal phosphoproteomics shows in vitro and in vivo vesicle-mediated signaling in response to EGF stimulation

The most significant advantage of our subcellular fractionation protocol is that it enables rapid and comprehensive proteome and phospho-proteome measurements, which provides an excellent basis for exploring the dynamic behavior of proteins in response to specific stimuli in a spatio-temporal manner. Dynamic cell signaling via Receptor Tyrosine Kinases (RTKs) represents an excellent model to study spatio-temporal protein regulation ^23,24^. Furthermore, epidermal growth factor receptor (EGFR) activation by ligand binding, auto-phosphorylation, recruitment of signaling adaptors and subsequent internalization into endocytic vesicles for either degradation or recycling is a great example of how spatiotemporal dynamics in cells are controlled by rapid phosphorylation events ^5^. To study EGFR signaling dynamics at the subcellular level, we stimulated HeLa cells with EGF and measured the change in the subcellular proteome and phospho-proteome at 5 different time points (*i.e.,* 0, 2, 8, 20 and 90 minutes upon EGF stimulation). We performed the experiment in biological quadruplicates. From the resulting 240 DIA runs, we were able to confidently quantify (in at least three out four replicates) 7142 unique proteins all covering different protein-coding genes (PCGs) and 11046 phosphorylation-sites (Fig. 3A). As expected, we found that EGFR was mainly purified in the plasma membrane fraction (fraction 4), although we also detected it in nuclear fraction 6, which has been previously described in literature ^25^ (Fig. 3B). Upon EGF stimulation, EGFR is rapidly auto-phosphorylated on its cytoplasmic tyrosine residues, which triggers the recruitment of adaptor proteins such as GRB2 ^26^, SHC1 ^27^ and CBL ^28^, transducers of downstream signaling. These series of events unleash the rapid internalization of the receptor into endosomes for further degradation ^5^. Since our approach does not separate plasma membrane from membrane-organelles (such as endosomes) we cannot directly follow the translocation of EGFR from the plasma membrane to early endosomes. However, we can clearly detect how the adaptor proteins GRB2, SHC1 and CBL, which are in all the cytosol in unstimulated cells, rapidly reduce their cytosolic presence upon 2 minutes EGF stimulation, as they are recruited to the EGFR containing membrane fraction (Fig. 3B). Most interestingly, we also traced the dynamic wave-like movement of these proteins across time, as they are shuttled back to the cytosol following the transient activation and degradation pattern of EGFR phospho-signaling after 20 minutes upon stimulation and recruitment by EGFR. This was measured directly by quantifying the dynamic phosphorylation of EGFR tyrosine residues, which follow the same dynamic pattern in the membrane fraction 4 as the adaptor proteins (Fig. 3C). These EGFR tyrosine phosphorylation sites act as docking sites for the SH2 domains in the adaptor proteins, which are subsequently phosphorylated, too. For example, we find that SHC1 is rapidly phosphorylated at tyrosine 427 after 2 minutes of EGF stimulation and that this phosphorylation is clearly observed in the membrane-bound fraction 4 (Fig. 3D). This observation indicates that SHC1 is activated upon contact with EGFR, therefore revealing that the EGFR phosphorylation of SHC1 is a direct consequence of subcellular translocation. Furthermore, we also detect signaling events downstream of EGFR, such as phosphorylation of the transcription factor JUN on serine 63 in its transactivation domain, which is important for its transcriptional activity showing how signaling from EGFR is transmitted into the nucleus (Fig. 3D).

**Figure 3:**
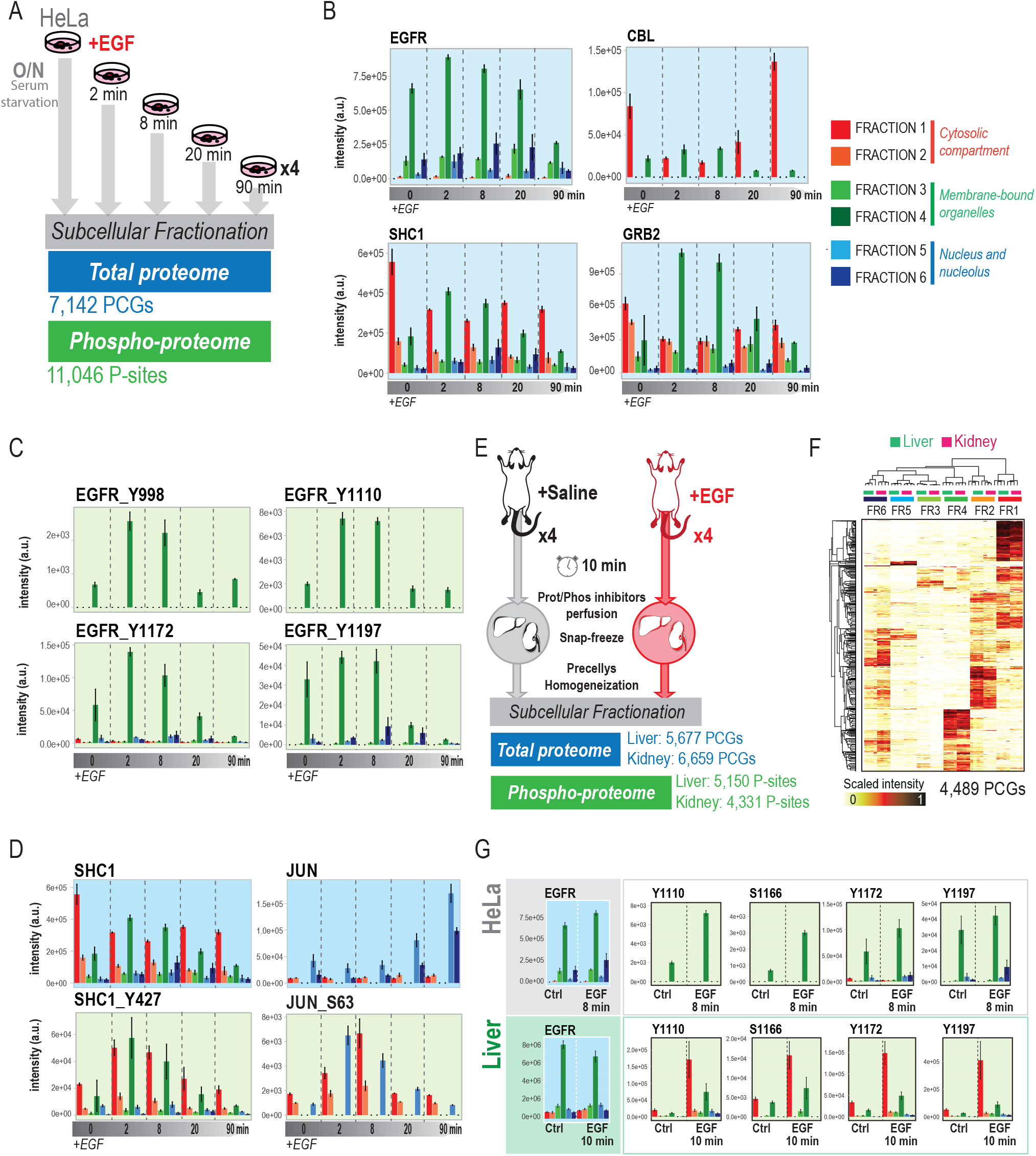
Spatio-temporal phosphoproteomics in response to EGF stimulation. (A) Experimental design and result overview with number of proteins and phosphorylation-sites (p-sites) obtained from the subcellular fractionation of HeLa cells treated with EGF at different time points. (B) Bar-plot of protein intensities across fractions and time points of EGFR and adaptor proteins CBL, SHC1 and GRB2. (C)Bar-plot of intensities across fractions and time points of EGFR tyrosine phosphorylation sites. (D) Bar-plot of protein intensities and phosphorylation sites across fractions and time points of SHC1 and JUN. (E) Experimental design and workflow of subcellular fractionation proteome and phospho-proteome analysis of EGF treatment in mice and a summary of identified proteins and phosphorylation sites in liver and kidney respectively. (F) Heatmap of scaled intensities per replicate, of four replicates, of the subcellular proteome of mouse liver (green) and kidney (pink), showing both protein and sample clustering. (G) Bar-plot of protein intensities and phosphorylation sites across fractions and time points of EGFR in HeLa and Liver. In all bar plots error bars indicate the standard deviation of the mean across replicates.

Although powerful as model systems for studying cell signaling, cell lines have certain limitations for *in vivo* extrapolation. Therefore, to extend the scope of our spatio-temporal proteomics approach, we applied the workflow to an *in vivo* system. For that purpose, we performed animal experiments, where two groups of four mice were injected intravenously with saline or EGF for 10 min, respectively, followed by 1.5 minutes perfusion with protease and phosphatase inhibitors ^29^. Whole livers and kidneys were subsequently explanted for spatio-temporal (phospho)-proteomics. The subcellular fractionation protocol was slightly altered for adaptation to organ tissues (see Methods). Briefly, tissues were homogenized in a saline buffer using the Precellys tissue homogenizer system. Then, tissue extracts were pelleted by centrifugation, and cleaned twice with a saline buffer before proceeding with the subcellular fractionation protocol. We were able to quantify 5677 and 6659 proteins across the six fractions in liver and kidney, respectively; as well as 5150 and 4331 phosphorylation-sites (Fig. 3E). We observed high reproducibility of the proteins purified in each fraction between tissues (Fig. 3F), and found that most of the subcellular compartments enriched per fraction reproduced what we previously observed for HeLa cells (Suppl. Fig. 4A). However, we noticed few but relevant differences mainly for the fraction in which lysosomal and extracellular matrix proteins were purified (Suppl. Fig. 4A-B). Interestingly, we also found that mitochondrial proteins showed a completely different profile in kidney versus liver, which might reflect the intrinsic differences in the morphology of this organelle between the two tissues ^30–32^.

To benchmark the spatio-temporal phospho-signaling observed *in vivo*, we extracted the spatio-temporal profiles of the cytoplasmic EGFR tyrosine phosphorylation-sites from liver phospho-proteome and compared their subcellular distribution to the corresponding data obtained from HeLa cells. As expected, we observed a significant increase in the intensity of the tyrosine phosphorylation-sites upon EGF stimulation in both liver and HeLa cells, but, interestingly, we found that in contrast to HeLa cells, these sites showed a dual distribution between cytoplasmic fraction (fraction 1) and membranous fraction (fraction 4) in liver cells (Fig. 3G). Considering the previously observed difference in the distribution of lysosomal markers between cell lines and tissues, we wondered whether this was also the case for endosomes. We therefore examined the distribution of protein markers for early (RAB5A, EEA1), late (RAB7A) and recycling (RAB6A, RAB8A) endosomes in the liver and kidney subcellular proteomes (Suppl. Fig. 4C) and, found that early endosomes markers were also co-purified in fraction 1 in the two tissues examined in here (Suppl. Fig. 4C). This observation can explain the difference in EGFR phosphorylation-sites observed, as activated EGFR is rapidly internalized into early endosomes, and consequently it is expected that the tyrosine phosphorylated EGFR sites would show the highest signal in the same fraction, where the early endosomes are found (Fig. 3G).

### Subcellular protein dynamics reveals ribosome accumulation upon osmotic stress

Once we proved the suitability of our approach to measure rapid protein translocation as a consequence of activated phospho-signaling networks, we decided to employ our methodology to identify subcellular relocation events triggered in response to cellular stress signaling. Specifically, we focused on the cellular response to hypertonic stress, which has already been described as a cause of protein translocation, as in neuronal cells ^33^. We treated U2OS cells for one hour with 500 mM sorbitol to induce hyperosmotic stress conditions. Moreover, to study the plasticity of the cells and their recovery from the osmotic stress, we washed out the sorbitol after the hypertonic stress event and collected cells in recovery after 30 minutes, 3 hours and 24 hours, respectively (Fig. 4A). Following our high-throughput mapping of the subcellular proteome and phospho-proteome, we were able to quantify 7588 proteins and 9462 phosphorylation-sites (Fig. 4A). To identify potential translocations as a consequence of osmotic shock, we calculated the ratios between the protein intensities after one hour of sorbitol treatment versus control conditions, for each fraction individually. The resulting ranked lists were used to perform Gene Set Enrichment Analysis (GSEA) ^34^ using the Cellular Component Gene Ontology (GO). Among the most significant GO terms enriched in this analysis, we found the “Cytosolic Large Ribosomal Subunit” gene set, which was significantly down-regulated in fraction 2 (Suppl. Fig. 5A). Interestingly, we observed that the same gene set in fraction 6 inversely mirrored the trend observed in fraction 2 (Suppl. Fig. 5A), indicating a potential switch between the two compartments. To further evaluate this observation, we measured the percentage of ribosomal proteins, for small and large subunits, respectively, in each subcellular fraction at all the time points (Fig. 4B). We observed that the increase of ribosomal proteins in fraction 6 was restricted to the 60S subunits, and not the 40S counterparts, indicating that the translocation is specific to large ribosomal subunit proteins and not the whole ribosome. In agreement with this, we also observed the decrease in 60S ribosomal content in fraction 2. Importantly, the ribosomal distribution reverted to original conditions, after 3 hours upon stress relief, suggesting a fast recovery from the stress conditions (Fig. 4B).

**Figure 4:**
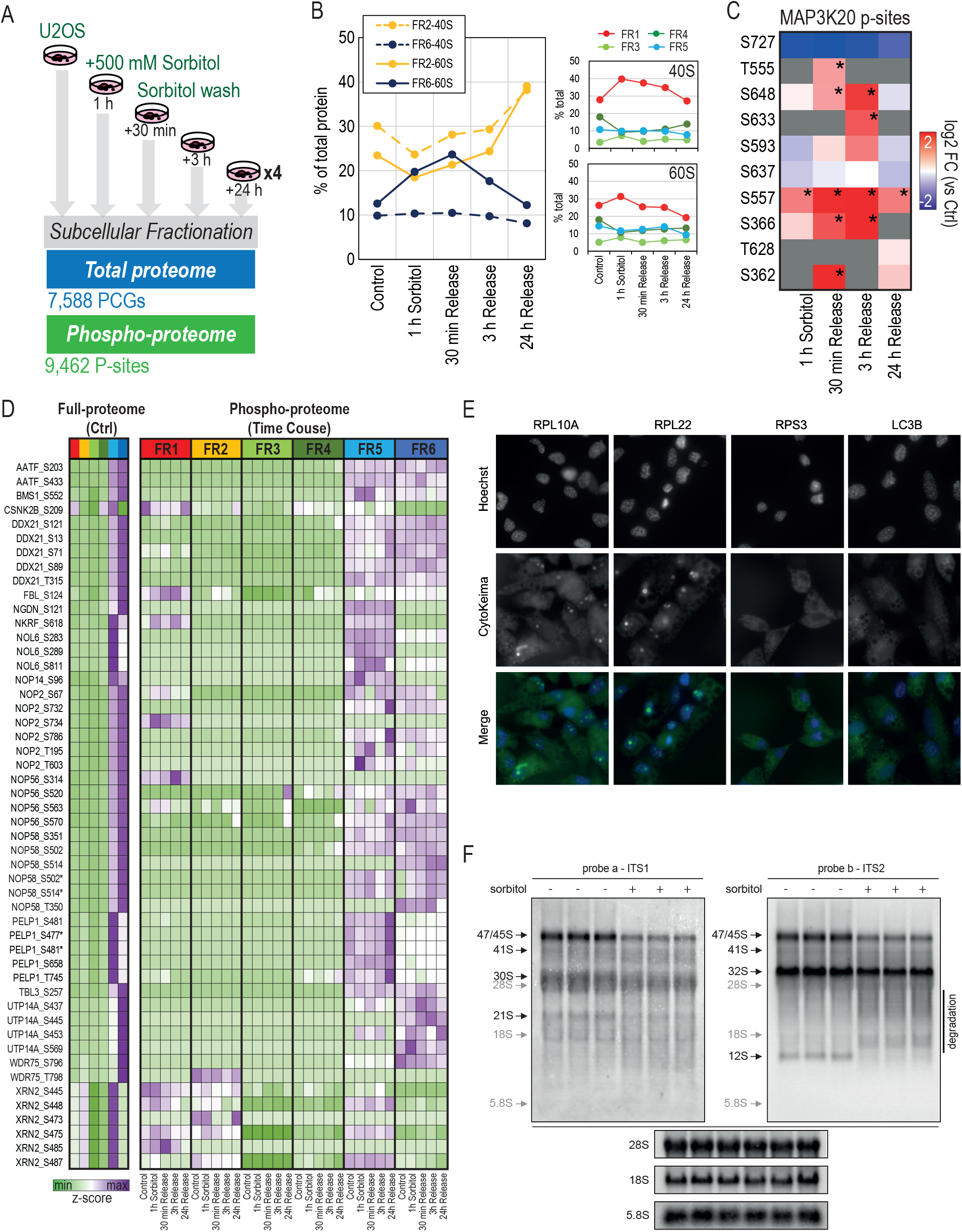
Subcellular protein dynamics during hyperosmotic shock. (A) Experimental design and result overview of subcellular fractionation of U2OS cells upon osmotic shock with sorbitol and posterior release. (B) Line-plot reflecting the percentage of total ribosomal protein (separately for 40S and 60S subunits) in each subcellular fraction at each given time point. (C) MAP3K20 phosphorylation sites regulation across time points. Intensity is depicted as log2 fold-change. Asterisks indicate statistical significance (moderated t-test, BH FDR q-value <0.05). (D) Heatmap of protein and phosphorylation site z-score intensities of ribosome assembly factors. Full-proteome intensity is only shown for initial/control conditions. (E) Representative images of TIG3 cells expressing mKeima-tagged RPL10A, RPL22, RPS3 or LC3B and treated with 500mM sorbitol for 3h and analyzed for pH neutral keima signal (CytoKeima). For RPL22 and LC3B n=4 and for RPL10A and RPS3 n=2. (F) Northern blots of whole cell RNA from biological replicates of U2OS cells treated with and without 500 mM sorbitol, probed with probe a targeting ITS1 (left) and probe b targeting ITS2 (right). Black arrows indicate rRNA processing intermediates (see Supplementary Figure 5E for a processing scheme) and gray arrows mark the migration of mature rRNA species.

The effects of hyperosmolarity can vary depending on intensity and duration of the treatment, but it mainly involves increased cellular toxicity, for which JNK and p38 MAP kinase signaling are key effectors ^35^, which are activated by upstream MAPKKKs (*i.e.:* MAP3K20 or ZAK). However, not much is known related to the downstream signaling elicited by hyperosmotic shock and posterior release. We observed a significant increase in p38-alpha activation loop phosphorylation (MAPK14 Y182) after 3 hours of release from osmotic stress, whereas phosphorylation of JNK1 (MAPK8 Y185) peaked after 30 minutes of release (Suppl. Fig. 5B). Interestingly, we observed rapid phosphorylation of p38 targets upon sorbitol treatment, such as STAT1-S727 (Suppl. Fig. 5B), whilst JNK targets had more delayed phosphorylation kinetics with for example JUN-S63 peaking after 3 hours of release (Suppl. Fig. 5B). Importantly, JNK and p38 stress signaling is known to be mediated by ZAK is response to ribotoxicity ^36–38^. We found a dual distribution of ZAK (MAP3K20) between the cytosol (fractions 1 and 2) and the nucleus (fraction 5), but where cytosolic ZAK in fraction 2 decreased significantly after 1 hour of hyperosmotic shock (Suppl. Fig 5C), that was not accompanied by a corresponding increase in its nuclear fraction. Conversely, several ZAK phosphorylation sites, identified in the cytosolic fraction, showed a distinct up-regulation trend that peaked between 30 minutes and 3 hours of stress relief (Fig. 4C). This might suggest a potential connection between ZAK activation and the ribosome translocation observed in our dataset. Moreover, the switch of 60S ribosomal proteins towards fraction 6 might indicate an accumulation of the 60S ribosome subunits in the nucleolus, where ribosome biogenesis and assembly occurs, which are highly complex and coordinated processes that require multiple factors ^39–42^. Altogether, our data suggest that osmotic shock induces some alteration in the ribosome biogenesis and assembly machinery leading to ribosome stalling in the nucleolus. To further investigate this, we extracted the subcellular proteome and phospho-proteome information of known ribosome assembly factors ^43^. As expected all of them were purified in nuclear and nucleolar fractions (Fig. 4D). We did not observe significant changes in their protein levels or subcellular localization upon osmotic shock. However, the phosphorylation status of several ribosome assembly factors, such as NOP58 or UTP14A, changed significantly (Fig 4D) indicating a functional modulation at this level due to the osmotic shock. To confirm our findings, we performed fluorescence microscopy in TIG-3 human fibroblast cell lines expressing Keima fluorescent markers for RPL10A, RPL22, RPS3 and LC3B. Fluorescent imaging revealed that upon hyperosmotic shock with 500 mM sorbitol, the large 60S ribosomal subunits (RPL10A and RPL22) showed significant accumulation in very condensed regions within the nucleus (Fig. 4E) likely representing nucleolar subcompartments. Most importantly, such foci were not observed either for the small (40S) subunit (RPS3) or for the control (LC3B) (Fig. 4E). Next, we evaluated whether the effect was purely sorbitol dependent by using decreasing concentrations of the osmolite, and observed that the 60S ribosomal protein accumulation is observed consistently with 250 mM sorbitol but not with 100 mM (Suppl. Fig. 5D). Collectively the data suggested that hyperosomotic shock triggers certain ribotoxicity that results in the accumulation of large ribosomal subunit proteins in the nucleolus, impeding its proper assembly and export to the cytosol. To pinpoint the origin of the defect leading to the observed accumulation, we performed a northern blot analysis in which two labeled probes targeting the ribosomal RNAs (rRNAs) required for small and large ribosomal assembly were analyzed. Probe “a” mapped the processing of rRNA components of the 40S ribosome (18S), while probe “b” mapped that for the 60S subunit (5.8S and 28S) (Suppl. Fig. 5E). The blot revealed that for the probe “b”, the band corresponding to 12S was missing and instead there was a smear indicating degradation of its precursor (Fig. 4F and Suppl. Fig 5F-G). Therefore, the degradation observed in the northern blot points to a defect on rRNA processing for the 60S subunit, which most likely explains the accumulation of the large ribosomal subunit proteins in the nucleolus.

### Muscle contraction in mice recapitulates ribosomal translocation observed in vitro upon osmotic stress

In addition to osmotic stress, there are other stimuli that elicit MAP kinase driven activation. One such example is the mechanical perturbation during muscle-fiber contraction, which like osmotic shock, activates JNK/p38 stress signaling ^44,45^. Therefore, we decided to validate whether the translocation of the ribosomal particles observed *in vitro* was also recapitulated *in vivo* after mechanical activation of the muscle. To do so, we performed animal experiments by exposing one of the lower hindlimbs of anaesthetized mice to a 10 minutes *in situ* contraction protocol, while the contralateral leg served as a resting control. This was followed by the immediate harvesting of tibialis anterior (TA) muscles from both legs, which were snap-frozen and cryo-pulverized before subcellular fractionation (Fig. 5A). Due to the minute amount of sample available and the well-known high dynamic range of the skeletal muscle proteome, the coverage of the subcellular proteome and phospho-proteome was limited to 3123 proteins and 1571 phosphorylation-sites. However, we were able to correctly identify 37 members of the 60S ribosomal subunits and 29 members of the 40S ribosomal subunits (~80% of total ribosomal proteins), which allowed us to evaluate location and dynamics of this organelle after mechanical contraction of the muscle. In fact, analogous to the osmotic stressed cells, we observed a similar dual distribution of the ribosomal subunit proteins between fraction 2 and fractions 5 and 6. Importantly, we confirmed that muscle stimulation also altered the distribution of the ribosomal subunit proteins, which significantly decreased in fraction 2, whilst increasing in fraction 6 (Fig. 5B), confirming the trend observed *in vitro*. Interestingly in muscle, we observed that the translocation was also true for the 40S ribosomal subunit proteins. Altogether, it indicates that mechanical stress recapitulates to certain extent the ribosomal translocation observed in vitro, but for the whole ribosomal complex, and not limited to 60S subunits.

**Figure 5:**
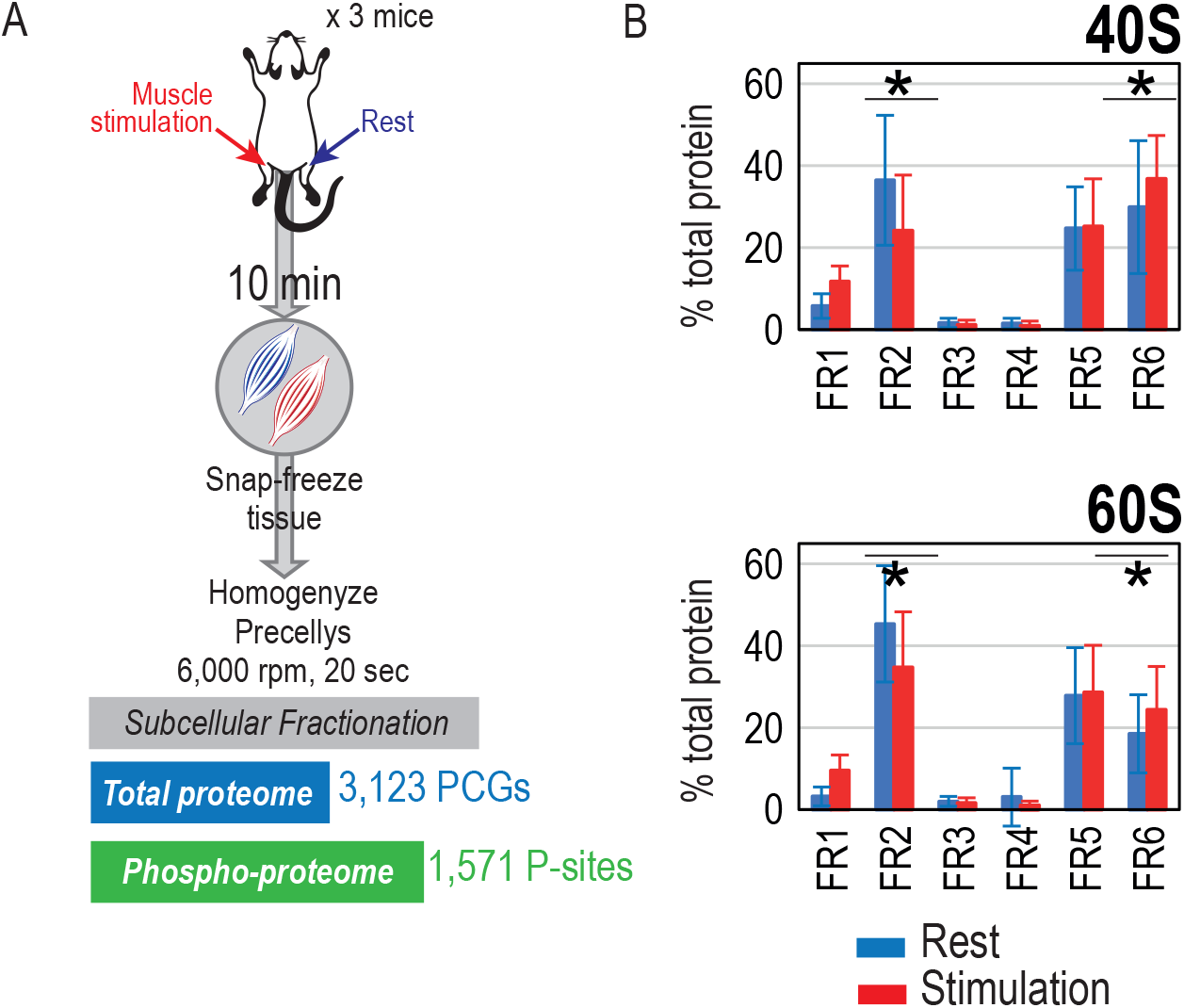
Muscle contraction in mice recapitulates ribosomal translocation. (A) Experimental design and workflow of subcellular fractionation proteome and phospho-proteome analysis of muscle contraction in mice. (B) Bar-plot of percentage of total ribosomal protein (40S top, 60S bottom) across fractions in resting conditions (blue) and after muscle contraction (red). Asterisks indicate p-value <0.01 in a paired two-sample t-test.

## Discussion

Here, we described a powerful approach to study spatio-temporal dynamics of the proteome and, to our knowledge for the first time, the phospho-proteome. Our workflow incorporates a simple and straightforward sequential cell fractionation protocol to profile six subcellular compartments with high reproducibility and scalability. For high-throughput analysis, we take advantage of recent developments in MS-based proteomics methods, such as the high-field asymmetric waveform ion mobility spectrometry (FAIMS) Pro interface and the Orbitrap Exploris 480 mass spectrometer to increase proteome coverage in short liquid chromatographic gradients for single-shot runs ^18^. Moreover, we also make use of a DIA approach, which circumvents certain limitations of more traditional DDA workflows for short LC-MS/MS runs, such as undersampling, dynamic range and reduced sensitivity. In addition, we employed a spectral library-free approach (*i.e.* directDIA in Spectronaut), which eliminates the necessity of spending MS-acquisition time to generate spectral libraries required for DIA-based (phospho)proteomics ^46^. Collectively, the optimized workflow minimizes the time required to obtain comprehensive maps of the subcellular proteome and phospho-proteome dynamics, which to date was a great limitation to current MS-based spatial proteomics approaches (Suppl. Table 1). Thanks to the reduction in time to map a whole proteome and its corresponding phospho-proteome of a sample, to just 5 hours of MS time, we were able to apply the workflow to multiple biological replicates, cell states, stimuli, and treatment time-points. This was not only to demonstrate the high reproducibility of the method, but also to employ it to study spatio-temporal dynamics in response to phospho-signaling networks activated by growth factors and stress.

While detergents with different solubilizing capacity have been used earlier for chemical subcellular fractionation, an important observation made here is that better partitioning can be achieved by also varying the ionic composition of the extraction buffers. Thus, cells that were permeabilized with a digitonin-based buffer 1 in a hypotonic buffer released a large number of proteins when resuspended in a detergent-free solution buffer 2 with 140mM NaCl (Fig. 2). Many of these proteins are annotated with the nucleus as the only location. As mentioned previously, we cannot exclude that the release occurred from mitotic cells with a disintegrated nuclear membrane. However, the absence of nuclear proteins in extracts obtained when cells were treated with Tween-20 in buffer 3 and dodecyl maltoside in a hypotonic buffer 4 suggests that ionic interactions are more important than membranes for retaining nuclear proteins. Successive use of Tween-20 and dodecyl maltoside (DDM) provides means to separate soluble proteins in cytoplasmic organelles from membrane proteins, while exposure of membrane-stripped nuclei to a detergent-free solution with 500 mM NaCl in buffer 5 yields a highly pure nuclear extract (Fig. 2). The remaining insoluble material is a mixture of nucleoli, cytoskeleton and poorly soluble membrane proteins that requires a 0.3% SDS-containing buffer 6 for solubilization.

On the other hand, we applied this chemical fractionation approach to show how EGFR adaptor proteins rearrange their subcellular distribution in response to EGF stimulation, after which they relocate from their free cytosolic form to the membrane fraction and EGFR-bound vesicles, and that this trend followed the phosphorylation activation of tyrosine residues in EGFR. However, this experiment reflected one major caveat of the current workflow, which is the impossibility to separate vesicles from plasma membrane proteins in the fourth fraction when applied to epithelial cell lines. This limitation impedes us to properly track the translocation of EGFR from the cell membrane to the endosome. One possible way to solve this could be to apply surface protein labeling ^47^ before the subcellular protocol is performed, such that surface proteins could be separately purified from fraction 4.

Importantly, we described the applicability of this workflow to rodent tissues opening the possibility to study spatio-temporal phospho-signaling regulation in *in-vivo* systems. However, we observed relevant differences in the cellular compartment profiles obtained from liver or kidney when compared to HeLa cells. This suggests that differences in the size, morphology and/or protein composition of organelles have a significant impact in their separation properties with this workflow. Interestingly, we observed a striking difference in mitochondrial protein distribution between liver and kidney. For the kidney, we found that most mitochondrial protein markers distributed very similarly to their HeLa cell profiles, whereas it seemed that the same mitochondrial proteins were purified in earlier fractions for the liver likely reflecting the morphological and phenotypic differences in hepatic cell mitochondria compared to those from other cell types. This might indicate the necessity to modify the protocol to adapt to tissue specificities if needed.

Finally, we have demonstrated how our approach can be applied to discover previously undescribed mechanisms of the cellular stress response. Although it was already known that hypertonicity can induce ribotoxic stress due to p38 activation, we have shown by using MS-based spatial proteomics that this ribotoxicity is impacting ribosome biogenesis and assembly resulting in accumulation of 60S subunits in the nucleolus, which could be explained by the defective rRNA processing machinery specific to 60S ribosomal subunit that we identified. Collectively, our *in vivo* and in vitro datasets represent a large resource of subcellular (phospho)-proteome dynamics. To make it available for other researchers in an easy accessible form, we have created a web-database SpatialProteoDynamics.github.io with a simple user interface that allows researchers to query our database of subcellular proteome and phospho-proteome dynamics for their protein of interest. Altogether, this manifests the usefulness of the methodology hereby presented for prospective studies of spatio-temporal regulation using MS-based proteomics.

## Ethics declarations

Mice experiments carried out in this study were performed according to the guidelines of the Danish Animal Welfare Act and the Directive 2010/63/EU for the protection of animals used for scientific purpose. The study was approved by the Institutional Animal Care and Use Committee of the University of Copenhagen and the Animal Experiments Inspectorate, Ministry of Environment and Food, Denmark (License number 2020-15-0201-00508 and 2019-15-0201-01659) and the Institutional Animal Care and Use committee of the University of Copenhagen (Project number P20-372 and P19-342).

## Competing interests

All authors declare no competing interests.

## Acknowledgements

The authors would like to thank Jørgen F.P. Wojtaszewski (University of Copenhagen, Denmark) for access to *in situ* muscle contraction equipment, and Miguel Garrido Rodrigo for his help setting up the web-based resource. Work at The Novo Nordisk Foundation Center for Protein Research (CPR) is funded in part by a generous donation from the Novo Nordisk Foundation (Grant number NNF14CC0001). The proteomics technology developments applied was part of a project that has received funding from the European Union’s Horizon 2020 research and innovation programme under grant agreements: MSmed-686547, EPIC-XS-823839, and ERC synergy grant 810057-HighResCells. The work was also supported by a research grant from The Novo Nordisk Foundation to A.L. (NNF20OC0059767).

## Author contributions

A.M.-V. designed, optimized and performed all experiments, analyzed the data and wrote the manuscript. D.B.B.-J. and S.S. helped perform the cell line experiments. K.S., A.M., T.T., and F.L.J. developed the cell fractionation protocol. F.L.J. developed the updated version of MetaMass and performed MetaMass-based analysis. E.T.V. and A.L. performed the in-vivo EGF stimulation experiments. E.K., S.B.-J., S.H.B. and L.B.F. performed fluorescent imaging experiments. R.K. performed *in situ* muscle contraction. N.K performed the rRNA processing experiment. J.V.O. designed the experiments, critically evaluated the results, analyzed the data, and wrote the manuscript. All authors read, edited, and approved the final version of the manuscript.

## Methods

### Buffer preparation for subcellular fractionation

The subcellular fractionation protocol requires the preparation of the following washing buffers: (i) washing solution A (30 mM Hepes pH 7.4; 15 mM NaCl, 2 mM MgCl2, 1 mM EDTA), (ii) washing solution AS (30 mM Hepes pH 7.4; 15 mM NaCl, 2 mM MgCl2, 1 mM EDTA, 350 mM sucrose) and (iii) washing solution AG (30 mM Hepes pH 7.4; 15 mM NaCl, 2 mM MgCl2, 1 mM EDTA, 20% glycerol). Just before starting the procedure, protease and phosphatase inhibitors were added to each buffer to get the following final concentrations: 1 mM TCEP, 1 mM NaF, 1 mM beta-glycerol phosphate and 5 mM of sodium orthovanadate. Additionally, one tablet of cOmplete™ Mini, EDTA-free Protease Inhibitor Cocktail was added to 10ml of the washing buffers.

### Cell culture and collection

U2OS and HeLa cells were grown in a P15 dish until 70-80% confluence. Cells were serum starved overnight then washed with PBS and harvested by trypsinization (1.5ml of trypsin). Trypsinized cells were resuspended in 8.5 ml ice-cold PBS containing 5 mM of Sodium-orthovanadate for a total volume of 10 ml and centrifuged for 3 minutes at 400 g. Cell pellets were washed twice with ice-cold PBS containing 5 mM of Sodium-orthovanadate. All subsequent steps were performed at 4°C.

### Subcellular fractionation

Cell pellets were resuspended in 540 μl of AS wash and 60 μl of 0.15% digitonin solution, which has been previously heated at 95°C for 5 minutes. Samples rotated on ice for 30 minutes and were spun down for 3 minutes at 500 g in a swing out rotor centrifuge. Supernatant was recovered and transferred to a clean tube labeled as Fraction 1. Cell pellets were washed twice with 1 ml of AS wash.

Cell pellets were resuspended in 540 μl of AS wash and 60 μl of 1.4 M NaCl. Samples rotated on ice for 15 minutes and were spun down for 3 minutes at 500 g in a swing out rotor centrifuge. Supernatant was recovered and transferred to a clean tube labeled as Fraction 2. Cell pellets were washed twice with 1 ml of AS wash.

Cell pellets were resuspended in 570 μl of AS wash and 30 μl of 10% Tween-20. Samples rotated on ice for 15 minutes and were spun down for 3 minutes at 500 g in a swing out rotor centrifuge. Supernatant was recovered and transferred to a clean tube labeled as Fraction 3. Cell pellets were washed twice with 1 ml of AS wash.

Cell pellets were resuspended in 540 μl of AG wash and 60 μl of 10% dodecyl maltoside. Samples rotated on ice for 15 minutes and were spun down for 3 minutes at 500 g in a swing out rotor centrifuge. Supernatant was recovered and transferred to a clean tube labeled as Fraction 4. Cell pellets were washed twice with 500 μl of AG wash.

Cell pellets were resuspended in 540 μl of A wash, 60 μl of 5 M NaCl and 1 μl of Benzonase^®^ Nuclease. Samples rotated on ice for 15 minutes and were spun down for 3 minutes at 2000 g in a fixed angle rotor centrifuge. Supernatant was recovered and transferred to a clean tube labeled as Fraction 5. Cell pellets were washed once with 500 μl of AG wash.

Cell pellets were resuspended in 522 μl of A wash, 60 μl of 1.4 M NaCl and 18 μl of 10% SDS. Samples were boiled for 10 minutes at 95°C. Vials containing this last fraction were labeled as Fraction 6.

All collected fractions were spun in a fixed angle rotor centrifuge for 5 minutes at 20,000 g and transferred to clean Eppendorf tubes. Samples were stored at −80°C for further analysis.

### Mice EGF stimulation and tissue collection

Littermate male C57BL/6JRj mice obtained from Janvier Labs (Le Genest-Saint-Isle, France) at six weeks of age were housed at the animal facility of the University of Copenhagen in individually ventilated static type II cages (Techniplast) with access to food (Altromin 1314, Altromin) and water *ad libitum* and a controlled temperature and relative humidity environment (22 ± 2°C and 55% ± 10%, respectively) with 12:12h dark:light cycle.

For EGF stimulation experiments, adult mice (eight weeks age, 22.1± 2.3 g weight) were assigned to two study groups of four mice each by simple randomization. Mice were anesthetized with 2% isoflurane in oxygen using a precision vaporizer (Leica Biosystems). In each group, four mice were administered sterile epidermal growth factor in isotonic saline (EGF, 100 μg/kg bodyweight) or isotonic saline intravenously in a single bolus dose into the inferior vena cava. 10 minutes post injection, the animals were perfused (1.5 minutes, 4.5 ml/min) with ice-cold isotonic saline containing protease inhibitors (Roche cOmplete™ Mini, EDTA-free Protease Inhibitor Cocktail) and phosphatase inhibitors (1 mM NaF, 1 mM beta-glycerol phosphate and 5 mM of sodium orthovanadate) using a syringe pump (Aladdin AL-1000, World precision instruments). Livers and right kidneys were quickly removed and snap frozen in liquid nitrogen. The total time from dosing to tissue collection was 12 minutes. For subcellular fractionation, only part of the median lobe of the liver was used.

### Mice muscle stimulation and tissue collection

For muscle stimulation, fed mice were anesthetized by an intraperitoneal injection of pentobarbital (10 mg/100 g body weight, diluted 1:10 in a 0.9% saline solution) and left to recover on a heating plate (30°C) for ~20 min. Subsequently, an electrode was placed on a single common peroneal nerve followed by 10 min *in situ* contraction of TA muscle. The contralateral leg served as a sham-operated resting control. The contraction protocol consisted of 0.5 seconds trains repeated every 1.5 seconds (frequency: 100 Hz; duration; 0.1 ms; voltage; 5V). TA muscle from both legs was removed immediately following euthanasia and snap frozen in liquid nitrogen.

Prior to subcellular fractionation, tissue samples were homogenized using a Precellys system with 3 beads (2.8 mm) for 20 seconds at 6,000 rpm in buffer A. After centrifugation at 2000 g, the supernatant was removed. Sample was washed twice in buffer A before starting the subcellular fractionation.

### Sample preparation for MS analysis

Subcellular fractions were denatured, reduced and alkylated with 0.3% SDS, 5 mM TCEP and 10 mM CAA during 10 minutes at 95°C. Afterwards, samples were digested overnight using the PAC protocol ^17^ implemented for the KingFisher robot as described previously ^18,48^. Briefly, samples were divided into two wells for digestion, so 300 μl of sample were digested in parallel. Protein aggregation was performed by adding 700 μl of 100% acetonitrile (ACN) and 50 μl of MagResyn-Amine beads. Samples were washed three times with 1 ml of 95% ACN and twice with 1 ml of 70% ethanol. Samples were digested in 300 μl of 50 mM ABC, LysC (in 1 to 500 ratio enzyme to protein) and trypsin (in 1 to 250 ratio enzyme to protein). Samples were acidified after digestion to final concentration of 1% trifluoroacetic acid (TFA). 20 μl of each sample were loaded directly into Evotips for full proteome analysis. Remaining sample was loaded onto Sep-Pak cartridges (C18 1 cc Vac Cartridge, 50 mg - Waters).

### Phospho-enrichment of subcellular fractions

In order to perform phospho-enrichment of each subcellular fraction, peptides previously loaded into Sep-Pak cartridges were eluted into the KingFisher plate using 75 μl of 80% ACN. 150 μl of loading buffer (80% ACN, 8% TFA and 1.6 M glycolic acid) was added to each sample. Phospho-enrichment was performed as described previously ^18,48^ using 10 μl of TiIMAC-HP beads (MagResyn, Resyn Bioscience). Eluted phosphopeptides were acidified with 10% TFA to pH <3 and loaded into Evotips for further MS analysis.

### LC-MSMS analysis

All samples were analyzed on the Evosep One system using an in-house packed 15 cm, 150 μm i.d. capillary column with 1.9 μm Reprosil-Pur C18 beads (Dr. Maisch, Ammerbuch, Germany) using the pre-programmed gradient for 60 samples per day. The column temperature was maintained at 60°C using an integrated column oven (PRSO-V1, Sonation, Biberach, Germany) and interfaced online with the Orbitrap Exploris 480 MS. When using FAIMS, spray voltage was set to 2.3 kV, otherwise it was set to 2kV, funnel RF level at 40, and heated capillary temperature at 275°C. For full-proteome analysis using DIA and FAIMS full MS resolutions were set to 120,000 at m/z 200 and full MS AGC target was 300% with an IT of 45 ms. Mass range was set to 350−1400. AGC target value for fragment spectra was set at 100%. 49 windows of 13.7 Da scanning from 361 to 1033 Da were used with an overlap of 1 Da. Resolution was set to 15,000 and IT to 22 ms and normalized collision energy was 27%. Compensation voltage for FAIMS was set to −45. For phospho-proteome analysis using DIA we employed 17 windows of 39.5 Da scanning from 472 to 1143 Da with 1 Da overlap. Resolution was set to 45,000 and IT to 86 ms. Normalized collision energy was set at 27%. All data were acquired in profile mode using positive polarity.

### Raw data processing

Full proteome and phospho-proteome subcellular fraction raw files were searched using Spectronaut (v14) with a library-free approach (directDIA) using either human database (Uniprot reference proteome 2019 release, 21074 entries) or mouse database (Uniprot reference proteome 2019 release, 22286 entries), supplemented with a database of common contaminants. Carbamylation of cysteines was set as a fixed modification, whereas oxidation of methionines and acetylation of protein N-termini were set as possible variable modifications. Additionally, for phospho-proteome analysis, and phosphorylation of serine, threonine and tyrosine were included as well. The maximum number of variable modifications per peptide was limited to 3. Only for phospho-proteome files, PTM localization cutoff was set as 0.75. Cross-run normalization was turned off. For protein quantification, major protein group aggregation method was changed to sum. Phospho-peptide quantification data was exported and collapsed to site information using the Perseus plugin described in Bekker-Jensen et al (see Code Availabitiy) ^46^. All remaining processing steps were performed in either Perseus (v1.6.5.0) or R (v3.6.2).

### Data Analysis

Data at protein and phospho-site level were processed using R (v3.6.2). For normalization, to remove experimental bias, as well as for imputation of missing values, each fraction was treated separately. Protein identifications without valid gene names were discarded. Data was log2 transformed and three valid values in at least one experimental group were required to preserve the protein or phospho-site. Most of the data analysis was performed using functions implemented in the Dapar package (v 1.18.3) ^49^ and following the data analysis pipeline of Prostar (v 1.18.4) ^49^. Normalization was performed using loess function from limma package ^50^. Imputation of missing values was performed in two steps: first partially observed values (*i.e.,* values missing within a condition in which there are valid quantitative values) were imputed using the KNN function (at protein level) and slsa function (at phospho-site level); secondly, values missing in an entire condition were imputed using the detQuant function from imp4p package. Finally, differential expressed protein and sites were calculated using limma (two-sided, BH FDR<5%), requiring at least three valid values in one of the two experimental conditions compared.

For all barplots, error bars indicate standard deviation of the mean of four replicates.

### MetaMass Analysis

The precision of partitioning of proteins in subcellular compartments was assessed using an updated version of the Excel-based application MetaMass^15^. A detailed user manual for the published version can be retrieved from the Nature Methods website (https://www.nature.com/articles/nmeth.3967#Sec12). Briefly, the input is a list of proteins assigned to groups by k-Means clustering of a dataset, or a combination of multiple datasets. The output includes assigned locations for each protein with scores for reliability (precision) and statistics for recall and precision for the proteins used as markers for subcellular compartments.

The user pastes the list of protein and assigned groups into the spreadsheet and clicks “buttons” to select among several built-in sets of markers for subcellular locations. Some sets correspond to single-location annotations from Uniprot, The Gene Ontology Consortium and the Compartments Database (all retrieved January 2021). Others are locations assigned in spatial proteomics studies ^7,22^. MetaMass assigns proteins within a given group to the same subcellular location based on the content of marker proteins. For example, if two proteins in the group are markers for cytosol, and none are markers for other locations, all proteins in that group are assigned to the cytosol with a precision of 1. If the group also contains a marker for e.g. nucleus, the proteins are still assigned to cytosol, but with a precision of 0.66. If all markers for a given location are assigned to the correct compartment, the precision and recall for that compartment is 1. *E.g.,* if a third is assigned to the wrong locations, the precision is 0.66.

MetaMass II has a wider range of marker sets than what was included in the published version. There are also data from recent spatial proteomics studies to facilitate meta-analysis. All datasets were normalized to the maximum signal value measured across the fractions. The classification is based on standard Excel functions, and the functions are displayed by selecting the cells. The worksheets are protected to prevent the user from accidentally editing cells that contain formulas. The password to unprotect the sheets to make modifications is “1”. The workbook has macros to automate the analysis. Experienced Excel users will know how to view and modify the codes.

### Northern blotting analysis of ribosomal RNA processing

Ten μg whole cell RNA from U2OS cells treated with and without 500 mM sorbitol for 3 hours was separated on a formaldehyde denaturing 1% agarose gel. Then transferred to a BrightStar-plus nylon membrane (Ambion) by capillary blotting, followed by UV-cross-linking. Probes (10 pmol each) were labeled with [γ-32-P]-ATP using T4 PNK (Thermo Fisher) and hybridized to the membrane one by one in hybridization buffer (4× Denhardts solution, 6× SSC, 0.1% SDS) overnight at 10°C below the Tm of the probe. The membrane was washed four times in washing buffer (3× SSC, 0.1% SDS), exposed to a Phosphor Imager screen, scanned by a Typhoon scanner (GE Healthcare) and analyzed using Fiji software. The membrane was stripped between each hybridization using boiling hot 0.1% SDS.

Probe a; TGGGTGTGCGGAGGGAAGC
Probe b; ACGCCGCCGGGTCTGCGCTTA
28S rRNA; GCTCCCGTCCACTCTCGAC
18S rRNA; CCAGACAAATCGCTCCACCAACTAAG
5.8S rRNA; CCGCAAGTGCGTTCGAAGTGT

### Generation of stable cell lines

mKeima fusion sequences were cloned into pLVX-TetOne-Puro backbone using In-Fusion or Gibson Assembly cloning kits according to manufacturer’s instructions. RPL10A, RPS3 and LC3B have an N-terminal mKeima tag, while RPL22 carries a C-terminal mKeima tag. Stable cell lines were generated by lentiviral transduction of TIG3 fibroblasts. To generate virus, HEK293 cells were transfected with these plasmids along with PAX8 and VSV-G expressing plasmids and the virus collected after 24h. After transduction, positive cells were selected with puromycin and subsequently FACS sorted for keima expression.

### Cell culture, treatments and Keima imaging

TIG-3 fibroblasts were grown in Dulbecco’s Modified Eagle Medium supplemented with 10% Fetal Bovine Serum and 1% PenStrep. Keima expression was induced for 72h before sorbitol treatment with 100ng/ml doxycyclin. Sorbitol was dissolved in media immediately before being added to the cells. Cells were imaged live in HBSS with 1:6000 Hoechst (H3570) using an ImageXpress Micro Confocal High-Content Imaging System (excitation 440nm, emission 620nm).

## Code availability

Custom R code used in the manuscript is available from the corresponding author on reasonable request. PTM collapse plugin requires Perseus and R (minimum version 3.6.0) to run and it is available at github.com/AlexHgO/Perseus_Plugin_Peptide_Collapse.

## Notes

https://SpatialProteoDynamics.github.io

### Competing Interest Statement

The authors have declared no competing interest.

