## Supplemental Information for "Spatial-proteomics reveal *in-vivo* phospho-signaling dynamics at subcellular resolution"

- Supplementary Table 1.
- Supplementary Figures 1-5.

**Supplementary Table 1.** Comparative table different MS-based subcellular fractionation protocols.

| Digitonin + Homogeneization+ Ultracentrifugation | hyperLOPIT: equilibrium density gradient ultracentrifugation | | | LOPIT-DC: differential centrifugation | Chemical fractionation | |
|---|---|---|---|---|---|---|
| Orre *et al* | Christoforou *et al* | Thul *et al* | Geladaki et al | Geladaki et al | Current Study | As shown in |
| 30609389 | 26754106 | 28495876 | 30659192 | 30659192 | - | PMID |
| A431<br>U251<br>MCF7<br>NCI-H322<br>HCC-827 | Mouse pluripotent stem cells (E14TG2a) | U2OS | U2OS | U2OS | HeLa<br>U2OS | Cell line(s) |
| 1x P15 (3x p10 A431) | ~100 million cells | ~300 million cells | ~280 milliion cells | ~70 million cells | 1x P15 (~20e6 cells) | Input |
| 5 fractions (FS1, FP1, FP2, FP3, FS2) | 8 fractions from density gradient (out of 20) + cytosol + chromatin | 20 fractions from density gradient + cytosol+chromatin | | 10 fractions | 6 fractions | # Subcellular Fractions |
| TMT 10-plex | TMT 10-plex | | | TMT 10-plex | - | Labeling |
| HiRIEF (2x72 fractions) | High-pH reverse phase chromatography (24 fractions) | | | High-pH reverse phase chromatography (18-22 fractions) | - | Offline Fractionation |
| DDA | DDA (SPS-MS3) | | | DDA (SPS-MS3) | DIA | Data acquisition method |
| 90 min | 120 min | | | 120 min | 21 min | LC Gradient Duration |
| 9 days | ~2-3 days (per TMT10plex experiment) | | | 1.5 days | 2.5 h | MS Acquisition time (per replicate) |

**Supplementary Figure 1**

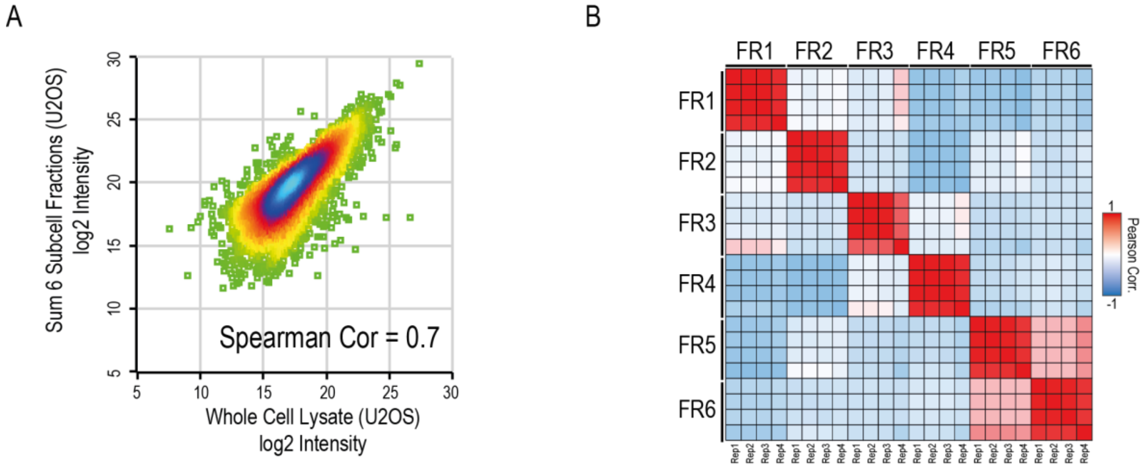


(A) Scatter-plot of whole cell lysate log2 intensities (as average of 4 replicates) against sum of log2 intensity of six subcellular fractions (as average of 4 replicates) obtained for U2OS cell line.

(B) Correlation plot showing Pearson correlation values between HeLa subcellular fractionation proteomics samples.

### Supplementary Figure 2

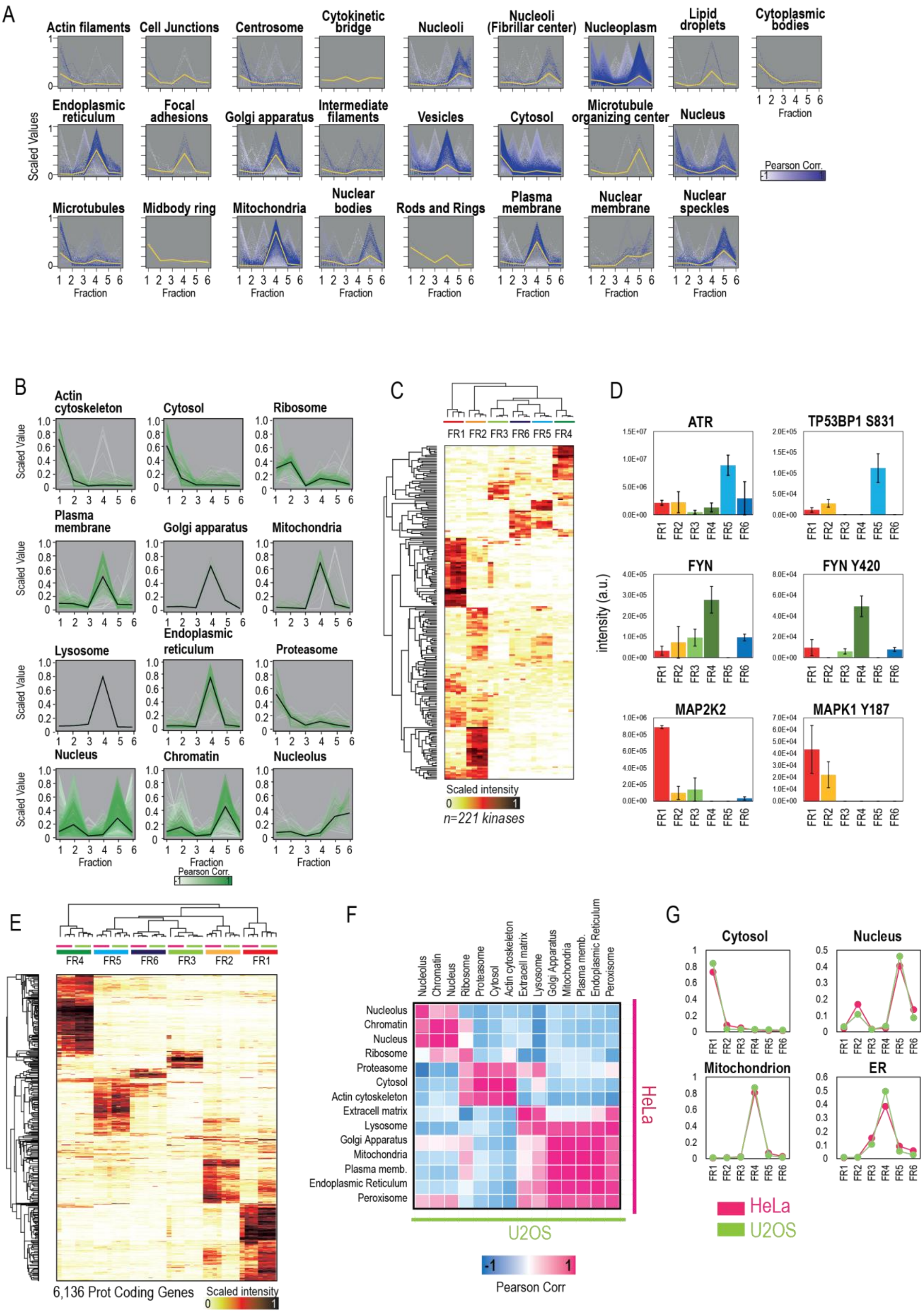

(A) Profile-plots of cell compartment markers obtained from The Cell Atlas (PMID: 28495876) in the subcellular proteome dataset. Scaled intensity across fractions is plotted for each independent replicate. Gradient of white to blue indicates Pearson correlation to the centroid of each distribution, which is highlighted as a yellow line.

(B) Profile-plots of cell compartment markers in the subcellular phospho-proteome HeLa dataset. Scaled intensity across fractions is plotted for each independent replicate. Gradient of white to green indicates Pearson correlation to the centroid of each distribution, which is highlighted as a black line.

(C) Heatmap of protein scaled intensities across fractions of the kinases present in the HeLa subcellular proteome dataset.

(D) Bar-plot of intensities across fractions in the HeLa subcellular fractionation datasets corresponding to protein kinases and representative phosphorylation substrates.

(E) Heatmap of protein scaled intensities across fractions of the kinases present in the HeLa (pink) and U2OS (green) subcellular proteome dataset.

(F) Correlation plot of the centroids of the distribution of cellular compartment markers between HeLa and U2OS datasets.

(G) Plot of centroids (measure as average per fraction of four replicates) of relevant cellular compartments in HeLa (pink) and U2OS (green).

**Supplementary Figure 3**

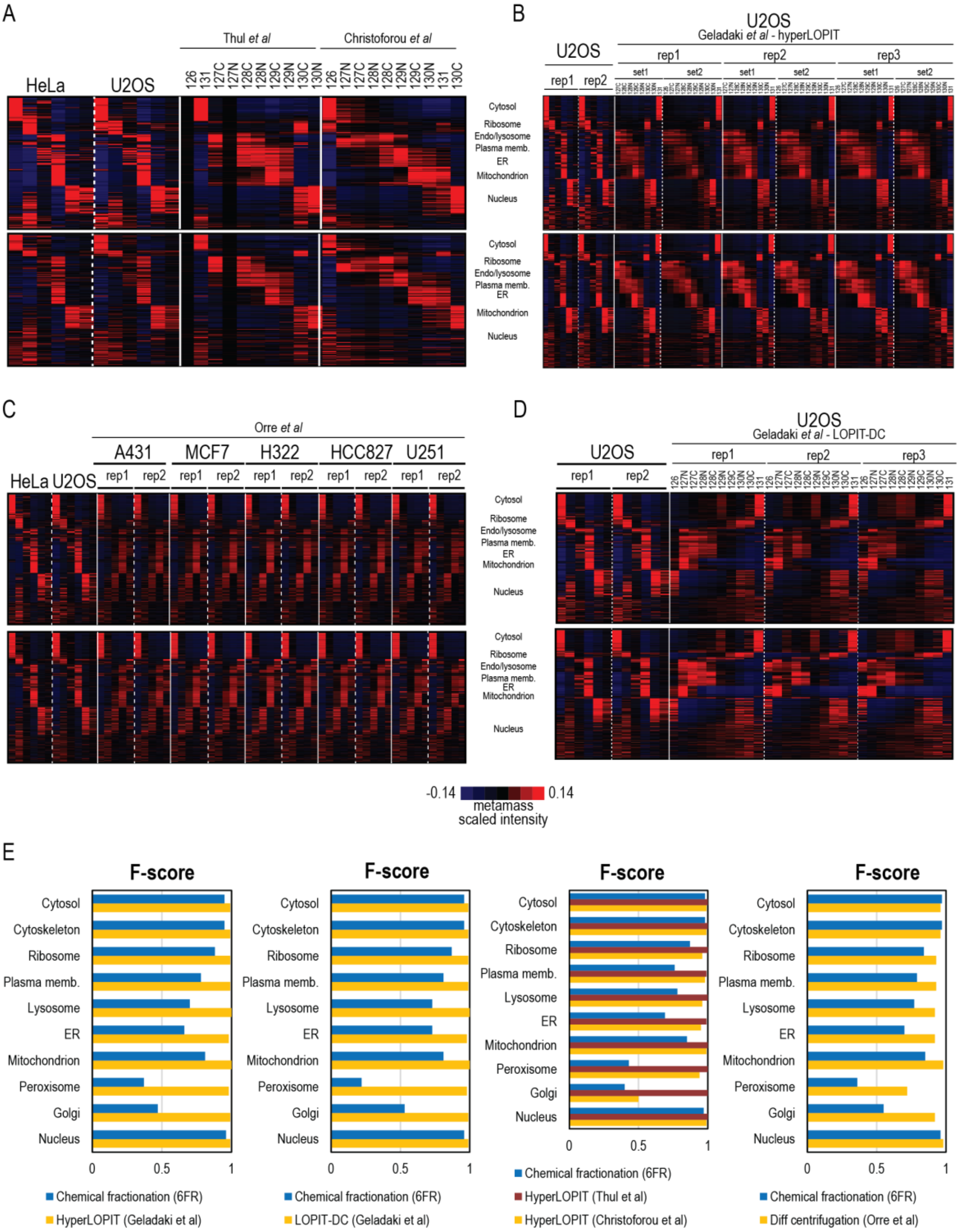


(A-D) Heatmaps showing protein distribution across fractions obtained from HeLa and/or U2OS using the present subcellular fractionation protocol and for other published studies. Proteins were classified and sorted using the Excel-based analysis tool MetaMass (Suppl. Table 4). All heatmaps were obtained after normalizing gene distribution and center samples by mean in Cluster 3.0, and plotted in TreeView. For all heatmaps, top heatmap corresponds to protein classification based on the data from this study, and bottom heatmap corresponds to protein classifications based on each corresponding published study.

(A) Comparison of protein distribution across fractions obtained from HeLa and/or U2OS using the present subcellular fractionation and HyperLOPIT subcellular fractionation method used in Christoforou *et al* and Thul *et al*.

(B) Comparison of protein distribution across fractions obtained from U2OS, either using the present subcellular fractionation and HyperLOPIT subcellular fractionation method used in Geladaki *et al*.

(C) Comparison of protein distribution across fractions obtained from HeLa and U2OS using the present subcellular fractionation and the different cell lines used in Orre *et al*.

(D) Comparison of protein distribution across fractions obtained from U2OS, either using the present subcellular fractionation or LOPIT-DC subcellular fractionation method used in Geladaki *et al*.

(E) F-score barplots for the protein assignment to organelles in the present study (blue) and different subcellular fractionation published studies (yellow and red).

**Supplementary Figure 4**

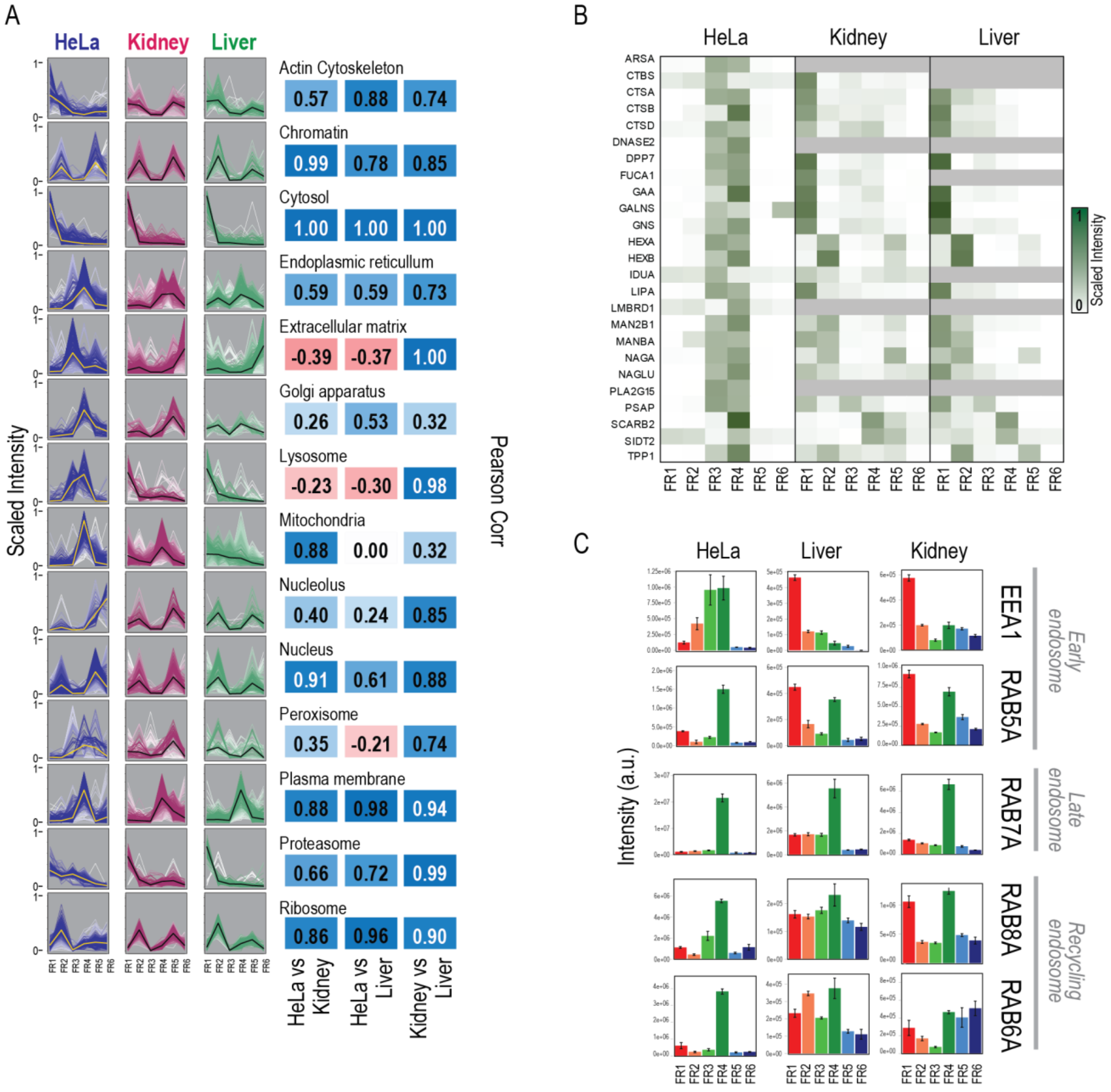


(A) Profile-plots of cell compartment markers in the subcellular proteome HeLa, Kidney and Liver datasets. Scaled intensity across fractions is plotted for each independent replicate. Gradient of color indicates pearson correlation to the centroid of each distribution, which is highlighted as a yellow/black line. Next to the plot, the Pearson correlation between samples of the centroids of each cell compartment is indicated.

(B) Heatmap of scaled intensities across fractions for lysosome markers in the HeLa, Kidney and Liver datasets.

(C) Bar-plot of protein intensities across fractions in the HeLa subcellular fractionation datasets corresponding to markers of early, late and recycling endosomes.

### Supplementary Figure 5

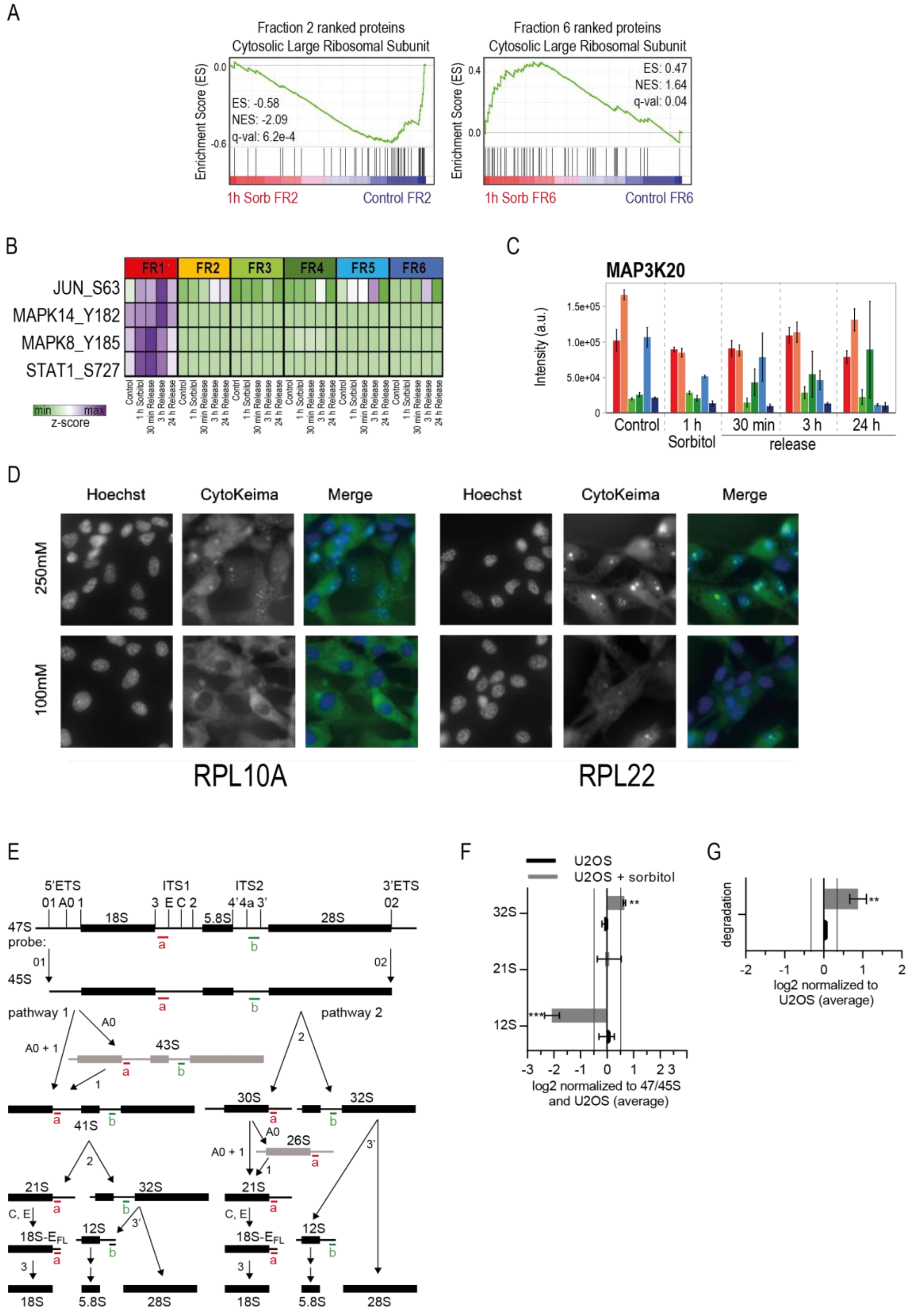

(A) GSEA plots for the GOCC term “Cytosolic Large Ribosomal Subunit" obtained from the protein ratios (1 hour Sorbitol vs Control) in fraction 2 and fraction 6.

(B) Heatmap of phosphorylation site z-score intensities of JNK and p38 signaling targets.

(C) Bar-plot of protein intensity across fractions and time points of MAP3K20.

(D) Representative images of TIG3 cells expressing mKeima-tagged RPL10A, RPL22, RPS3 or LC3B and treated with 500mM sorbitol for 3h and analyzed for pH neutral keima signal (CytoKeima). For RPL22 and LC3B n=4 and for RPL1Aa and RPS3 n=2.

(E) Scheme of the human rRNA processing intermediates with annotated processing sites and a simplified outline of the two main processing pathways with short-lived precursors in grey. The position of probe a and b used in Figure 4F are in red and green, respectively.

(F) Quantification of a subset of rRNA intermediates from northern blots in Figure 4F expressed as log2 fold change, internally normalized to 47/45S and the average of the three lanes containing RNA from control cells.

(G) Quantification of the area marked “degradation” in the right northern blot in Figure 4F expressed as log2 fold change and normalized to the average of the three lanes containing RNA from control cells.
